# Fluorescence resonance energy transfer and protein-induced fluorescence enhancement as synergetic multi-scale molecular rulers

**DOI:** 10.1101/047779

**Authors:** Evelyn Ploetz, Eitan Lerner, Florence Husada, Martin Roelfs, SangYoon Chung, Johannes Hohlbein, Shimon Weiss, Thorben Cordes

## Abstract

Advanced microscopy methods allow obtaining information on (dynamic) conformational changes in biomolecules via measuring a single molecular distance in the structure. It is, however, extremely challenging to capture the full depth of a three-dimensional biochemical state, binding-related structural changes or conformational cross-talk in multi-protein complexes using one-dimensional assays. In this paper we address this fundamental problem by extending the standard molecular ruler based on Förster resonance energy transfer (FRET) into a two-dimensional assay via its combination with protein-induced fluorescence enhancement (PIFE). We show that donor brightness (*via* PIFE) and energy transfer efficiency (*via* FRET) can simultaneously report on e.g., the conformational state of dsDNA following its interaction with unlabelled proteins (*BamHI, EcoRV*, T7 DNA polymerase gp5/trx). The PIFE-FRET assay uses established labelling protocols and single molecule fluorescence detection schemes (alternating-laser excitation, ALEX). Besides quantitative studies of PIFE and FRET ruler characteristics, we outline possible applications of ALEX-based PIFE-FRET for single-molecule studies with diffusing and immobilized molecules. Finally, we study transcription initiation and scrunching of *E. coli* RNA-polymerase with PIFE-FRET and provide direct evidence for the physical presence and vicinity of the polymerase that causes structural changes and scrunching of the transcriptional DNA bubble.

## INTRODUCTION

Advanced microscopy methods have become powerful tools for structural studies of biomolecules. These methods can complement classical biochemical and biophysical techniques^1,2^, but most importantly emerged as key player in understanding structural dynamics^3,4^. The underlying biophysical concept is straight forward: Construct a onedimensional molecular ruler, in which the biochemical state of the system can be read out as a distance-related measure. Such a molecular ruler often uses a photophysical property such as fluorophore brightness or lifetime to provide information on the structure of biomolecules in real time.^5^

A classic example for such a molecular ruler is Förster-type resonance energy transfer (FRET)^5^, which allows achieving structural information with a spatial resolution in the nanometre range and (sub)millisecond temporal resolution.^6–10^. However, other photophysical effects such as photo-induced electron transfer (PET)^11–14^ or protein-induced fluorescence enhancement (PIFE)^15–33^ can be used for similar purposes. Since the fluorescent signal can be read out with high time-resolution, even fast conformational changes^34–40^, as well as interactions between biomolecules, can be mapped in physiologically relevant environments *in vitro* ^41,42^ and *in vivo*^43,44^ with a sensitivity allowing to address individual molecules. These molecular rulers suffer from limitations such as their restricted distance ranges and the need for labelling with fluorescent dyes. Most importantly, in the assessment of a three-dimensional (dynamic) structure, the largest limitation is embedded in the information accessible to these methods, which at best follow a single distance to capture a complex structural state.

This restriction prohibits monitoring an essential feature of biological processes. While single distances can be read out with high spatio-temporal-resolution, it remains challenging to simultaneously observe conformational changes in different protein parts, map these structural changes as either a result of protein binding or due to intrinsic dynamics^45^ and observe how multi-subunit proteins coordinate conformational changes between different domains. To tackle these problems, multiple distances need to be monitored simultaneously (as read-out for the respective biochemical states). Multicolour approaches have been employed, e.g., FRET assays with more than two different fluorescent labels, but are not routinely used for complex biological systems due to difficulties in terms of physical instrumentation, molecular biology or labelling chemistry.^46–50^

Notably, the PIFE method provides a molecular ruler for sensing the proximity (<3 nm) of a DNA-bound protein to a Cy3-labeled DNA base. The method relies on the well-established fluorescence enhancement of Cy3 in high viscosity or in sterically restricted environments^51–58^. The enhancement of fluorescence is based on the competition between Cy3 photoisomerization from Trans (bright) to a 90° intermediate (dark) and deexcitation from Trans^52,57,58^. Although PIFE has been initially used in ensemble experiments^20–26^, its use following Cy3 fluorescence at the single molecule level (smPIFE) allowed the identification different protein-interaction modes using immobilized molecules^17,27–33^. PIFE has been combined with other techniques including bulk Cy3 PIFE and quenching^59^, smPIFE and single molecule nanomanipulation with flow^33^, and the combination of PIFE and FRET both in ensemble^20,21,24^ and immobilized single molecule experiments^30,60–62^. Although the combination of PIFE and FRET at the single molecule level was performed for immobilized assays, with its advantages and disadvantages, that method combination was not yet addressed for single molecule measurements in the freely-diffusing mode.

In this paper, we combine two fluorescence-related effects into one powerful assay that we dub ALEX-based PIFE-FRET. It allows observing changes in biochemical structure and interactions by following two distances with two different distance dynamic ranges. Strikingly, ALEX-based PIFE-FRET requires labelling with only two fluorescent dyes, i.e., similar to FRET. Its enhanced information content is provided by use of additional photophysical parameters, which are extracted via advanced data analysis procedures using single-molecule fluorescence detection and alternating-laser excitation. In detail, we utilize the stoichiometry parameter, S, as a measure for PIFE-effects (which may report on the vicinity of a protein bound to a DNA duplex), while FRET reports on the distance between fluorophores (which may report on the global conformation that a DNA duplex adopts upon protein binding). PIFE-FRET does not necessarily require surface-immobilized biomolecular complexes and hence obviates the use of other complex techniques such as ABEL-trap ^63,64^, feedback loop tracking ^65^ or use of microfluidic devices^66^.

To successfully construct and use a two-dimensional ruler thereby introducing PIFE for solution-based single-molecule experiments, we provide a framework for data analysis routine to allow simultaneous and quantitative read-out of two different photophysical parameters: donor brightness (PIFE) and energy transfer efficiency (FRET). In proof-of-concept experiments we study different oligonucleotide samples containing the environmentally sensitive donor Cy3 (PIFE fluorophore; FRET donor) in comparison to the fluorescence signals of the less environment-sensitive FRET donor Cy3B, combined with the acceptor ATTO647N. We show that PIFE-FRET enables the detection of the interaction between unlabelled proteins and doubly labelled diffusing dsDNA via changes in brightness ratio S (termed stoichiometry) using μs-ALEX^67^. We further investigated the spatial sensitivity of PIFE-FRET for binding of DNA-binding enzymes with respect to the PIFE-and FRET-ruler aspects. The modulation of donor-brightness due to PIFE (and hence the Förster radius) preserves the FRET-distance information after careful data evaluation. Finally, we study DNA scrunching in transcription initiation of *E. coli* RNA-polymerase (*RNAP*). Using PIFE-FRET, we provide the first direct evidence for simultaneous structural changes in the formed DNA bubble (→ DNA open bubble formation and DNA scrunching *via* FRET) accompanied by the physical presence of the RNAP protein (→ the vicinity of the bound RNAP to the promoter DNA *via* PIFE). Interestingly, we were able to identify an extrusion of the nontemplate strand of the transcription bubble out of the active site, recently identified through classic footprinting experiments^68^, yet without the use of chemical cross-linking reagents. The extrusion of the nontemplate strand was coupled to specific conformational changes in the bubble, caused by DNA scrunching. Lastly, we outline possible applications of PIFE-FRET both in studies with diffusing and immobilized molecules indicating the full potential of the technique for mechanistic investigations of biomolecular interactions.

## RESULTS

**The principles of ALEX-based PIFE-FRET**. The aim of the PIFE-FRET assay is to monitor two distances simultaneously with different dynamic ranges in complexes between proteins or nucleic acids and proteins. In the assay, we label two complementary ssDNA strands with donor (D) and acceptor (A) fluorophores both encoding distinct DNA binding sites after annealing (Fig. 1A). The brightness of the Cy3 donor fluorophore is increased upon binding of proteins in close proximity (Fig. 1C, PIFE), hence directly reports on the distance R1 between fluorophore and the surface of the bound protein (Fig. 1A).

**Figure 1.**
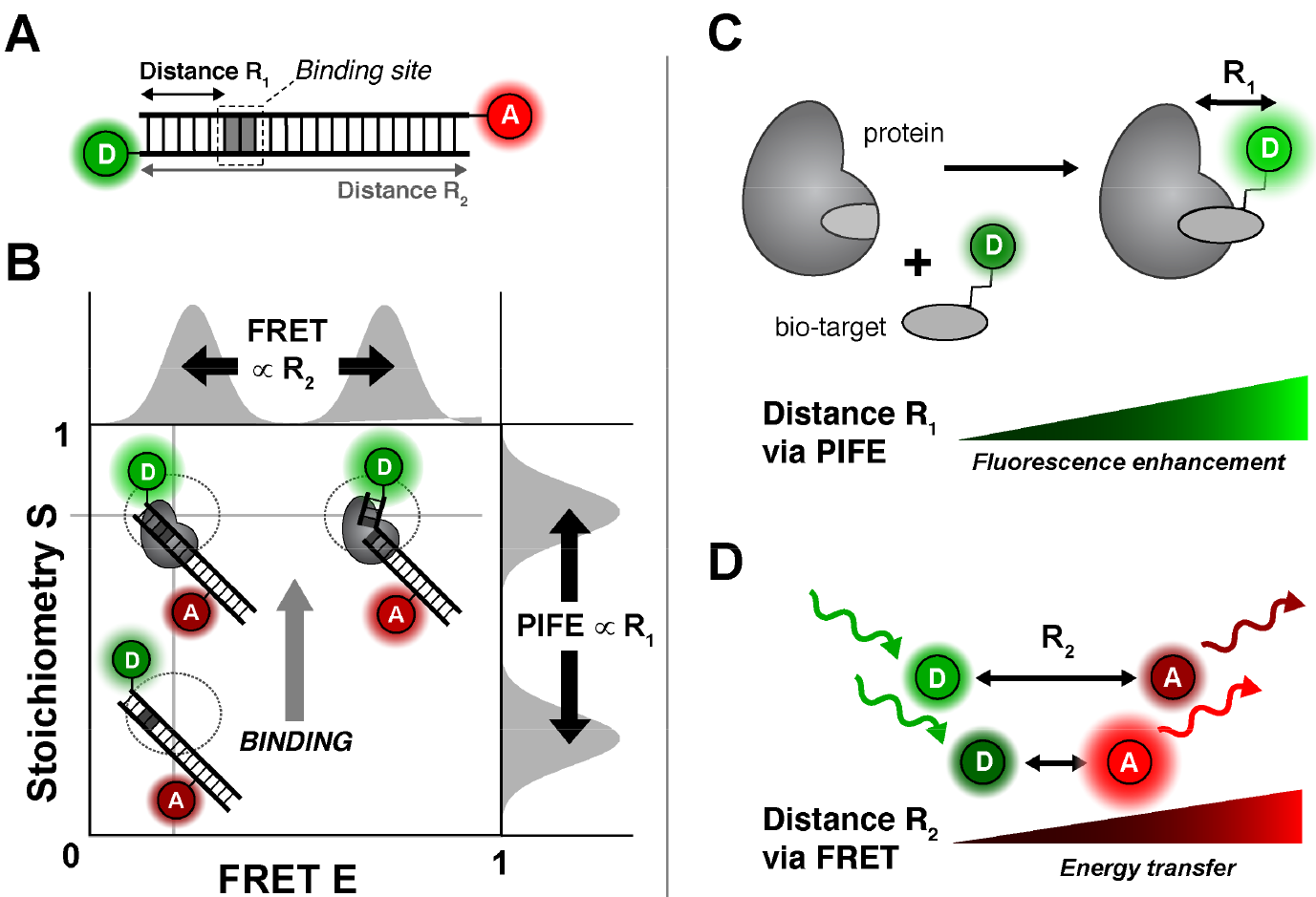
**Working principle of ALEX-based PIFE-FRET**. (A) Possible labelling scheme for PIFE-FRET experiments: a DNA template is labelled with FRET donor D and acceptor A and contains a binding site for a protein. (B) Read out of PIFE and FRET distances via ALEX: E-S-histogram depicts that changes of R_2_ can be monitored via FRET efficiency E, whereas distance R_1_ between donor and protein are determined by changes in stoichiometry S. (C/D) Photophysical effects reporting on distances R_1_ and R_2_: (C) PIFE reports on distance R_1_ between donor and the surface of a bound protein via fluorophore brightness. (D) FRET between D and A reports on distance R_2_.

The conformation of dsDNA is monitored by FRET (Fig. 1D, FRET) via changes in the interprobe distance R_2_ between D and A (Fig. 1A). A donor fluorophore that fulfils the requirements of the PIFE-FRET assay is the green cyanine dye Cy3 (Fig. S1), in which photoisomerization competes directly with FRET on excitation energy in a dye microenvironment-dependent manner. Finally, an experimental technique is required for simultaneously reporting on both photophysical parameters. For this we suggest to use alternating laser excitation (ALEX), which reports on PIFE effects via the ratiometric parameters, stoichiometry, S, and FRET efficiency, E (Fig. 1B). Both parameters of ALEX can be used for mapping out PIFE-FRET since S is defined as brightness ratio between the donor and acceptor fluorophore (Eqn. 6) and is hence sensitive to PIFE (as long as the brightness of the acceptor fluorophore is stable and unchanging) while E compares only donor-excitation-based intensities to derive the FRET efficiency (Eqn. 3–4).

While the proposed assay and its implementation in ALEX is as a straightforward combination to increase the information content of PIFE and FRET, the interdependency of photophysical properties represents a fundamental hurdle and will therefore be addressed carefully. Since the green donor fluorophore is integral part of both rulers, both PIFE and FRET compete directly via non-radiative de-excitation of the donor (Fig. 2).

**Figure 2.**
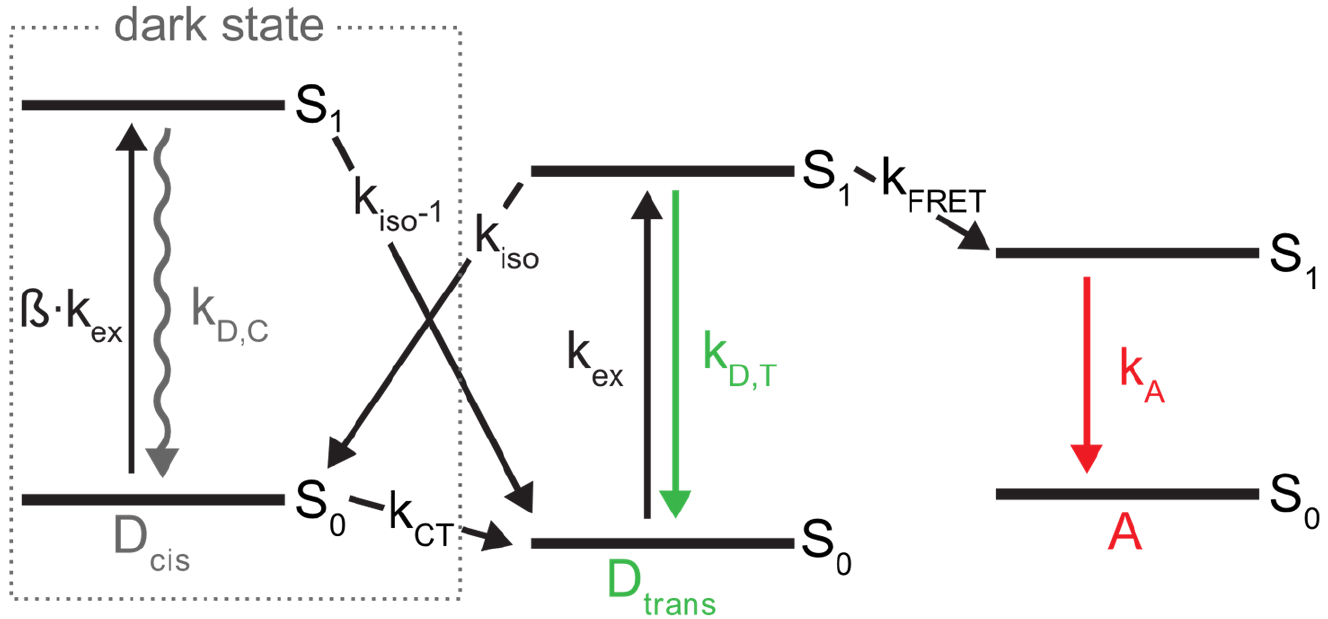
**Jablonski diagram of Cy3 in the presence of a FRET acceptor**. After excitation (*k_ex_*) to the excited trans isomer (D_trans_), three competing pathways deplete the excited state S1: (a) *k_D,T_* which is the sum of radiative and non-radiative decay rates from S_1_ to S_0_ resulting in fluorescence emission; (b) trans to cis photo-isomerization with *k_iso_* resulting in the formation of the non-fluorescent *cis* isomer D_cis_ of Cy3; (c) Förster-type energy transfer *k_FRET_* to an acceptor fluorophore A. The rates for Cy3 cis/trans isomerization *k_iso_* and *k_iso_*-1 are sensitive to the environment and are modulated by PIFE. ß accounts for differences in the extinction coefficients of cis and trans isomer at the excitation wavelength.

The quantum yield of Cy3 is environmentally sensitive, i.e., the PIFE effect alters both the non-radiative cis/trans isomerization (k_iso_) and the FRET parameters (Förster radius, R_0_ and hence the rate of energy transfer, k_FRET_) without changes in the distance between the donor and the acceptor. We therefore first describe a photophysical model as well as present a procedure for data evaluation in order to decouple both effects and determine PIFE (distance R_1_) and FRET (distance R_2_) in one experiment using a ratiometric approach rather than fluorescence lifetimes of the donor fluorophore^69,70^.

**Photophysics of PIFE-FRET**. The fluorophore Cy3 and similar cyanine fluorophores (DyLight 547, DyLight 647^71^, Cy5, Alexa Fluor 647) have been employed for PIFE as they all share the property of excited-state cis/trans isomerization^51,53–57^ that leads to dependence of their fluorescence quantum yield with respect to the local environment.^53^ This dependence includes the specific location of dye-attachment (for DNA 5’/3’ vs. internal labelling) as well as three-dimensional structure (single-vs. double-stranded DNA).^53,56,72^ The effect can be used to monitor interactions between an unlabeled protein and a fluorophore-labelled macromolecule (e.g., a dsDNA duplex Fig. 1C). Restricting the rotational freedom of the fluorophore by applying steric hindrance/restriction^73,74^, and also possibly specific interactions with select protein residues, the interacting protein causes a delay in photoisomerization to dark excited-state isomers^52,58^, which, in turn, increases fluorescence intensity without chromatic changes (Fig. S1). This effect allows the observation of complex formation between e.g., a Cy3-labelled oligonucleotide and a DNA-binding protein.^16^ In special scenarios (i.e., where suitable calibration is possible) PIFE can even serve as a quantitative ruler to determine fluorophore-protein proximity at base-pair resolution in a distance range of up to 3 nm that is inaccessible to smFRET^69,70^.

The underlying photophysical principle of PIFE was recently studied in detail by Levitus and co-workers^75^: The trans isomer of Cy3 is thermally stable, while the lifetime of non-fluorescent, ground-state *cis*-Cy3 is found in the μs range (Fig. 2, k_ct_).^76^ The deexcitation of isolated excited-state *trans*-Cy3 occurs primarily via two pathways, i.e., *photoisomerization* from *trans* (bright) to other (dark) excited-state isomers with rate k_iso_ and fluorescence after deexcitation from the *trans* isomer with rate k_D,T_ (Fig. 2).^77^ Under typical single-molecule conditions, i.e., continuous green excitation, both isomers *cis/trans* are populated during a fixed time interval with a population distribution determined by the microenvironment, which mainly alters rates k_iso_ and k_iso’_ (Fig. 2). Both rates depend on the energetic barriers between the isomers in excited-state (and the rotational mobilities/diffusivities for crossing them) between the corresponding S1-minima and a 90°-twisted geometry of Cy3 (Fig. S1).^75^ In principle, all other rates (except k_iso_ and k_iso’_) remain constant upon a change of the local environment, provided that these changes do not invoke different microenvironment quenching (e.g. stacking of Cy3 terminally labeled DNA, which shows changes in fluorescence^78^). The PIFE effect of Cy3 corresponds to a change in the population distribution between *trans* and the other (*cis* and the 90°-twisted intermediate) isomers, since it is mostly the trans isomer that contributes to a fluorescent signal (Fig. 2; Fluorescence quantum yield of *cis* isomer is negligible^79–81^ and the 90°-twisted intermediate isomer has a non-planar geometry, therefore it is nonfluorescent). Consequently, the mean fluorescence quantum yield, QY, of Cy3 (not its spectrum, Fig. S2) varies with environmental polarity, steric hindrance/restriction, (micro)viscosity and temperature^51,53–57^ and is thus sensitive to the steric restriction from adjacent binding of biomolecules such as proteins or DNA. It was shown experimentally that the isomerization rate constants are altered mainly in their pre-exponential factors due to a stronger dependence on diffusivity^51^. It was also shown that photoisomerization can be fully blocked by creating structural rigidity as in the derivative Cy3B (Fig. S1), leading to strongly reduced environmental sensitivity and increased brightness.^53,82,83^ Cy3B hence serves as a fluorophore that can emulate the maximal PIFE-effect since photoisomerization is fully prohibited/abolished in this molecule. When PIFE is combined with FRET, the donor-excited state is altered by both changes in k_iso_ and k_FRET_, and both rulers are directly dependent of each other (Fig. 2), however a change in k_iso_ is independent of the D-A distance R2. Therefore, after applying corrections to account for the changes in the QY of the donor, one can elucidate both PIFE and FRET information.

**Characterization of PIFE-signatures in ALEX experiments**. In order to understand the experimental signature of PIFE in FRET assays, we compared the spectroscopic properties of Cy3 and Cy3B in bulk and single-molecule experiments. Bulk fluorescence measurements of Cy3(B) on dsDNA were performed in the presence of increasing concentrations of iodide. These experiments show how both Cy3 and Cy3B are collisionally quenched at the same iodide concentration scale (Figs. S1C/D). However, Stern-Volmer plots show that the fluorescence QY of Cy3B depends linearly on Iodide concentrations, whereas Cy3 shows a nonlinear relation that is characteristic for a system with a sub-populations of fluorophores accessible and inaccessible to the iodide quencher (Fig. S1E).^84^ As expected, the relative QY of Cy3 further increased with increasing glycerol concentrations, whereas the relative QY of Cy3B is almost unaffected and remained constant under similar conditions (Figs. S1F/G). Increasing bulk viscosity of Cy3 affected fluorescence intensity with only negligible chromatic changes. These experiments suggest that PIFE effects indeed mainly influence the QY of Cy3 and hence the intensity of associated fluorescent signals.

To directly read out PIFE effects in confocal single-molecule fluorescence microscopy, we utilized ALEX (Fig. S2A and Methods Section) for studies of fluorescently-labelled dsDNA.^8,67,85,86^ Here, either Cy3 or Cy3B were combined with the FRET-acceptor ATTO647N. In ALEX histograms, the stoichiometry, S, allows to sort different molecular species according to their labelling: donor-only labelled dsDNA (S>0.8), acceptor-only labelled dsDNA (S<0.2) and donor-acceptor-labelled dsDNA 0.8>S>0.2 (Supplementary Fig. 2B). Since we are mostly interested in the properties of the species with both donor and acceptor labelling, we represent the final data set using a dual-colour burst search showing S over the proximity ratio E_PR_. S at this stage is denoted S(E_PR_) in Figure 3; the data was corrected for background and spectral cross talk^49^ after an all-photon burst search.

At a separation of 40 bp between donor and acceptor fluorophores (>10 nm), the proximity ratio is zero due and forms a prominent population in the ALEX histogram at intermediate peak S-values of ~0.3 (Fig. 3A, Cy3). Values for E_PR_ and S(E_PR_) were derived from a two-dimensional global Gaussian fit (see Material and Methods) represented as a black circle at full-width-half-maximum (FWHM).

**Figure 3.**
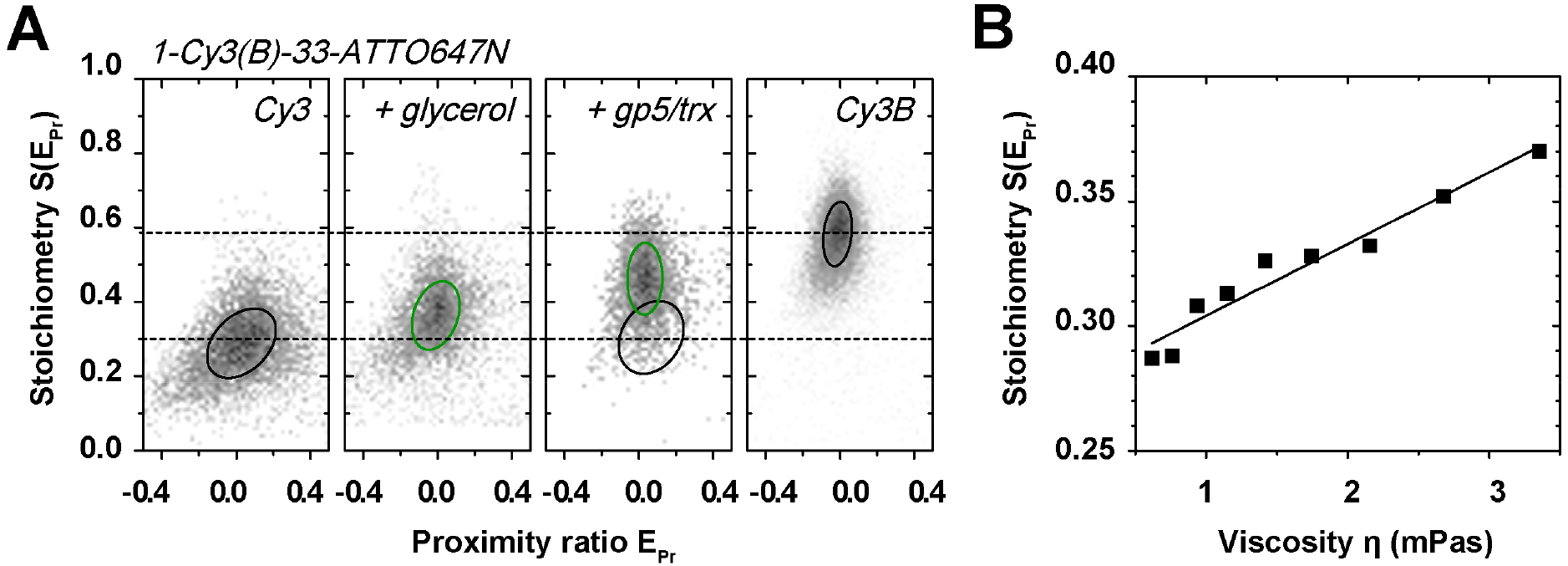
**Observing the PIFE effect in ALEX histograms**. (A) Cy3-ATTO647N, Cy3-ATTO647N with 40% glycerol, Cy3-ATTO647N in the presence of 30 nM T7 polymerase gp5/trx and Cy3B-ATTO647N. 2D Gaussian fitting was applied to characterize the observed populations; black/green circles mark the FWHM of each distribution in the presence (green) and absence (black) of brightness changes of the donor Cy3. (B) S(E_PR_) as a function of viscosity^87^ η using different concentrations of glycerol. The presented data is based on 45-mer dsDNA with large donor-acceptor separation of 33 bp (see Fig. S4A for oligonucleotide sequences and labelling positions). Relative differences of stoichiometry were found to be independent of laser power.

Addition of glycerol can emulate PIFE due to increase in bulk viscosity, which drives an increase in the mean fluorescence quantum yield of the Cy3 donor (Fig. 3A/S1F). As expected for such an increase, the donor-based photon streams DD and DA increase (data not shown), which hence increases S(E_PR_) upon addition of glycerol to the imaging buffer (Fig. 3A, Cy3 + glycerol). We find a linear increase suggesting that the PIFE-effect might indeed serve as a molecular ruler (Fig. 3B). A similar effect, as for increasing viscosity through the addition of glycerol, is observed upon steric hindrance caused by non-specific binding of T7 polymerase gp5/trx^88–90^ to dsDNA in close proximity to the Cy3 donor (Figure S3A). In Figure 3A a prominent PIFE population is seen at elevated S(E_PR_)-values when using concentrations of gp5/trx >50 nM (Figs. S3B/C). Similar PIFE-effects with a donor-enhanced QY, i.e., increase in S(E_PR_), are found for Alexa Fluor 555 (Fig. S3D) or with acceptor-enhanced quantum yield for ATTO647N and Cy5 (Figs. S3E/F). The herein presented results indicate that PIFE-FRET provides the capability to binding of unlabeled protein.

In ALEX-based PIFE-FRET, ideally only one fluorophore should be influenced by protein binding while the other should have an constant fluorescence signal for normalization of the PIFE-sensitive signal. This is best fulfilled in case of a small protein footprint not allowing interaction with the acceptor or use of acceptor fluorophore that is micro-environment insensitive. In our hands, only TMR and Cy3B show little to no PIFE effect (Fig. S3) while Cy3, Alexa555, Cy5 and ATTO647N are all influenced by protein binding.

To determine the maximally achievable PIFE-effect for Cy3, we investigated the dsDNA labelled with Cy3B/ATTO647N and compared the S-value to Cy3/ATTO647N (Fig. 3A). The ~4.1-fold increase in brightness between Cy3 and Cy3B poses a practical upper limit as seen through the reported, maximal PIFE-effect of ~2.7-fold brightness increase for binding of the restriction enzyme *BamHI* on dsDNA.^15–19^ A comparison of S(E_PR_) values between both samples hence emulates the maximal PIFE effect for ALEX experiments. In the absence of FRET, we find a range of S(E_PR_) from 0.29 (Cy3) to 0.59 for Cy3B (Fig. 3A).

**Spatial sensitivity of the PIFE ruler in ALEX**. Next, we investigated the spatial sensitivity of the PIFE ruler (distance R_1_) in ALEX experiments. By using a similar experimental scheme as introduced by Hwang et al.^16^, we designed different dsDNAs with a sequence to accommodate specific binding of the restriction enzyme *BamHI* (Fig. S5) at different R_1_ distances from the donor Cy3(B) (Fig. 4A). The donor-acceptor R_2_ distance was kept constant at 40 bp separation resulting in zero peak FRET efficiency (not to be mistaken with a donor-only species), while the binding site for the restriction enzyme was varied from 1,2,3,5 and up to 7 bp relative to the donor binding position (Fig. 4A). See Materials and Methods for the precise labelling scheme of the used dsDNAs. Restriction enzymes provide an excellent model system to study PIFE-FRET due to the well-characterized biochemical and structural behaviour^91–93^; both enzymes have been crystalized on dsDNA in the presence of calcium and bind as a homo-dimer (Fig. S5; pdb code: 2BAM). *BamHI* forms a stable, pre-reactive complex on dsDNA containing the palindromic sequence GGATCC without changing the conformation of the dsDNA. In a modified assay, we explored EcoRV binding to a GATATC-site. In contrast to *BamHI*, binding of EcoRV results in a tightly bound dsDNA conformation bent by 50°.

Figure 4B shows experimental data of *BamHI* binding to different positions on DNA revealing the ruler-character of ALEX-based PIFE (Fig. 4B/S6). Upon addition of 500 nM *BamHI* ^94^, we observed two sub-populations in S: the isolated Cy3-containing DNA (S(E_PR_) ~ 0.3) and a new PIFE-related population, i.e., *BamHI* bound to DNA at higher S(E_PR_) values (Fig. 4B). Optimal concentration for the *BamHI* ruler was determined by monitoring PIFE with different *BamHI* concentrations (Fig. S6); 1-Cy3-40- ATTO647N(1bp) showed a K*d* of ~ 40 nM. It should be noted, however, that the affinity between *BamHI* and the respective DNA varies for different positions of the *BamHI* binding site (see amplitudes of the PIFE-species in Figs. 4C/S6E). The control experiment of BamHI binding in close proximity (1 bp) to Cy3B shows a nearly unaltered peak stoichiometry upon binding to the dsDNA, which reports on a small decrease of Cy3B-intensity (Fig. S10A).

**Figure 4.**
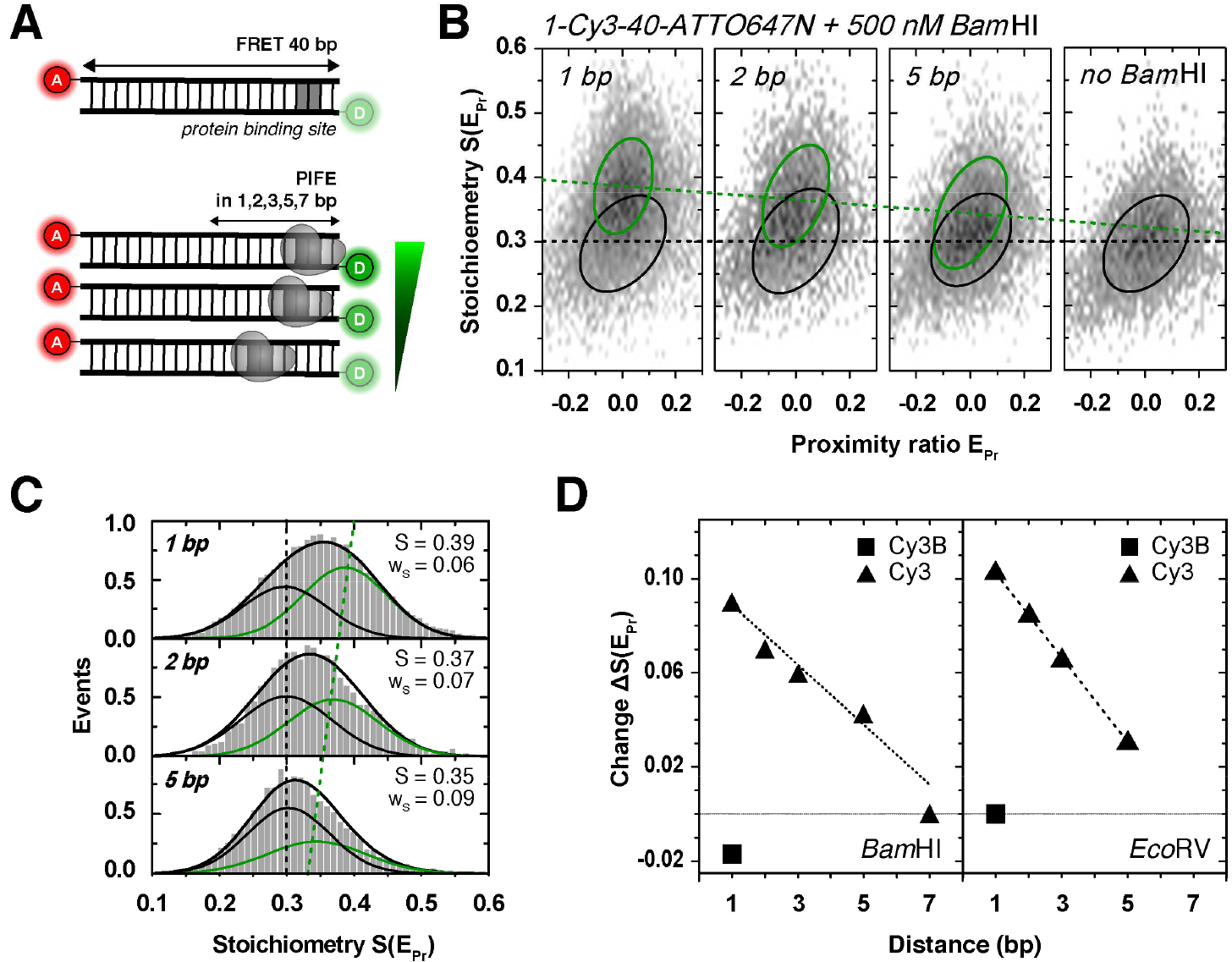
**The PIFE ruler in ALEX microscopy**. (A) Schematic of a dsDNA template containing a protein binding site of the restriction enzymes *BamHI* and *EcoRV* positioned in R_1_ = = 1,2,3,5 and 7 bp distance from donor fluorophore Cy3. The acceptor ATTO647N is positioned at the opposite 5’-end in R_2_ = 40 bp distance. (B) 2D histograms of 40 bp-dsDNA: free DNA (black circle) and DNA bound to 500 nM *BamHI* with PIFE (green, circle) in 1,2 and 5 bp distance. 2D Gaussian fitting was applied to characterize the observed populations; black/green circles mark the FWHM of each distribution. (C) *BamHI*-bound 40 bp-dsDNA: one dimensional projections of S(E_PR_) data and 2D Gaussian fits and their centre positions (free form, black; protein-bound form, green). (D) Absolute changes in S(E_Pr_) as function of R_1_ between Cy3 and *BamHI* (left) or *EcoRV* (right). Note that the control experiment with Cy3B (square) showed only little change of S(E_Pr_) compared to Cy3 (triangle) even at close proximity of Cy3B and protein of 1 bp.

The distance dependence of the PIFE-ruler reported here is approximately linear with similar dependency as reported^16^, i.e., 1-7 bp (Fig. 4D). We stress out, however, that it is not yet clear whether there is a general (linear) PIFE distance dependence, but published literature suggests that each system has to be evaluated independently. As an example, Nguyen et al. have lately identified an exponential PIFE ruler for the interaction of human replication protein A with ssDNA^61^ (nevertheless, this dependence is not decoupled from the possible flexibility effects of the Cy3-labeled ssDNA). The actual position of the Cy3 fluorophore (terminal vs. internal on the DNA) was found to have only a small influence on the observed S changes (Fig. S7), consistent with the fact that also stacked Cy3 can indeed photoisomerize^78^. The absolute values of PIFE-readout S, however, differ for comparable laser powers between internal and terminal labelling. Hence the donor brightness and also the PIFE effect changes with the labelling position on DNA. This effect should be considered when designing PIFE-FRET assays. When performing identical experiments as for *BamHI* (Fig. 4) with a different restriction enzyme (*EcoRV*, Fig. 4D/S8), we found a steeper distance dependence with an even more pronounced PIFE-effect. These results reveal the universal nature of the PIFE-ruler, but also that its specific quantitative properties require careful characterization for each biomolecular system - not only when used in PIFE-FRET but any other bulk or single-molecule assay.

**Calibration of the two rulers: R0-correction for Cy3-PIFE in the presence of FRET**. In the preceding section we have shown that the PIFE-ruler has a clear signature in ALEX experiments that renders it a useful tool for mechanistic biomolecular studies (Figs. 1C and 4). Since Cy3-PIFE is based on a competition of the radiative fluorescence transition k_D,T_ and the (non-radiative) isomerizsation k_iso_, its combination with FRET is complicated by the fact that energy transfer also depletes the donor excited state via k_FRET_ (Fig. 2). S depends only on the donor-excitation based fluorescence intensities that are altered by PIFE when using an environmentally insensitive acceptor. Different peak E values can indicate (i) real distance changes between the donor and acceptor or (ii) changes in R_0_ caused by altered donor QY for PIFE. A direct comparison of FRET efficiencies and related molecular distances for species with donor Cy3 and Cy3B with PIFE is impossible due to their different Förster radii R_0_ - a situation that is similar to comparing the FRET efficiencies of donor-fluorophores Cy3 and Cy3B in combination with identical acceptor fluorophores. When comparing a DNA-based ladder of Cy3(B)/ATTO647N with R_2_ separations of 8, 13, 18, 23, 28 and 33 bp (Fig. S4A), the problem becomes evident. We use corrected data E_PR_ / S(E_PR_) to read out brightness differences (Figs. 5A/D left panel; PIFE as an indicator distance R_1_). A direct comparison of the S-shift indicative of PIFE decreases directly with increasing FRET (Fig. 5D) - indicating the competition between FRET and PIFE^52^. This data suggests that the dynamic range of the PIFE-FRET assay regarding distance R_1_ (the range between S for Cy3 and for Cy3B, for a given E value) is optimal at low FRET efficiencies ^52^. Accurate FRET values^49^ are required to obtain distances on the FRET axis, i.e., distance R_2_. A comparison of identical dsDNA having either Cy3 or Cy3B as a donor with 13 and 23 bp R_2_ separations from the acceptor reveal significant differences in their peak accurate FRET E values (Fig. 5B). These differences of fluorophore pairs with identical interprobe distance reflect the shorter R_0_ of Cy3-ATTO647N (R_0_ = 5.1 nm^95^) as compared to Cy3B-ATTO647N (R_0_ = 6.2 nm^96^).

**Figure 5.**
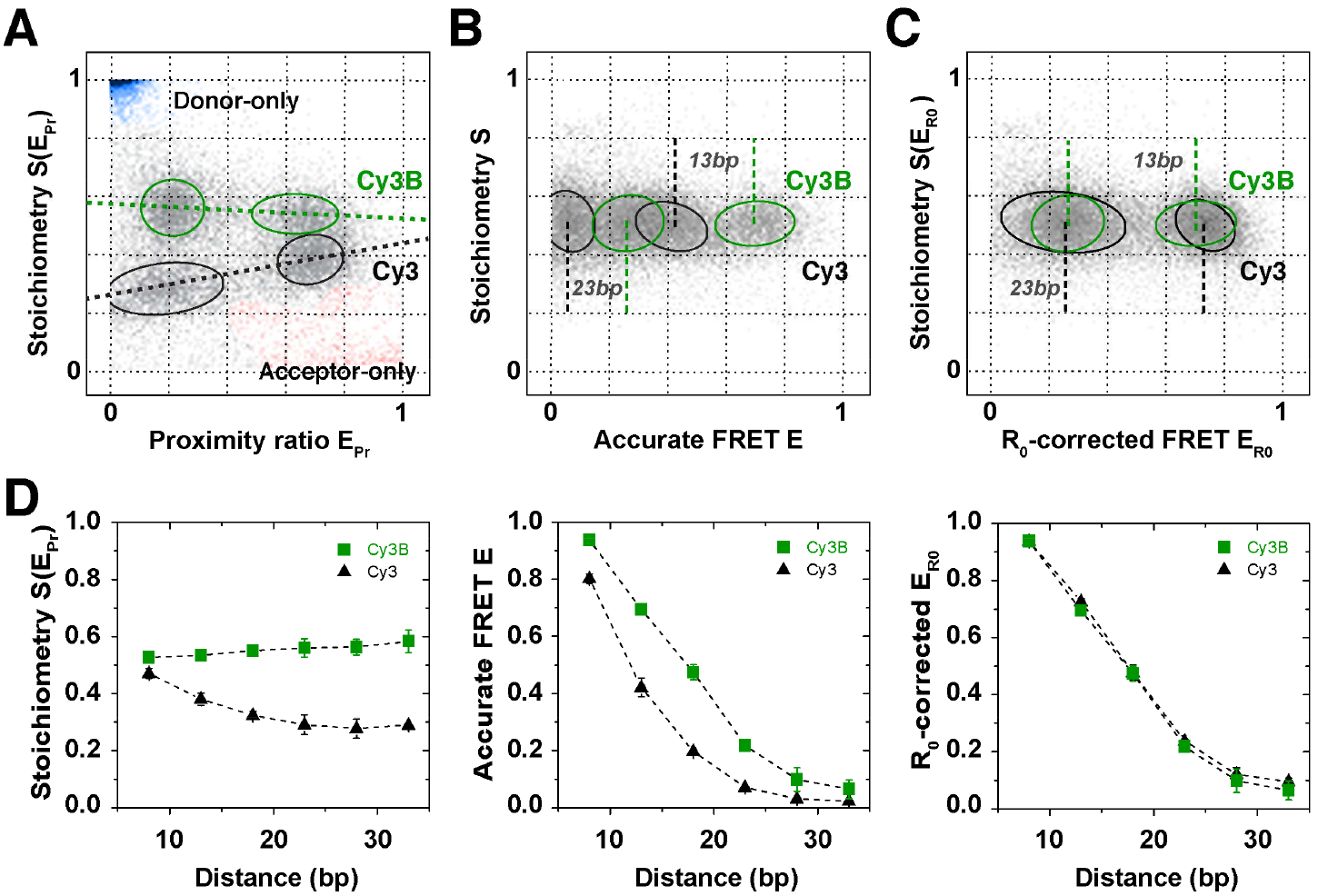
**Validation of the PIFE-FRET correction procedure in ALEX (A) Data correction process, to obtain R_0_-corrected 2D-histograms**. Four dsDNAs with identical sequence (Fig. S4A) are labelled with two different FRET pairs: Cy3(B)/ATTO647N and mixed together. The donor is attached at the 5’- end; the difference in brightness between Cy3 (black) and Cy3B (green) separates the four populations into two groups according to S(E_Pr_). The acceptor fluorophore ATTO647N is positioned on the complementary DNA strand in 13 and 23 bp distance; the two distances are deciphered via two different E_Pr_ values per subgroup. (B) By correcting each fluorophore pair with its corresponding gamma factor γCy3 or γCy3B, accurate FRET values E for each population are obtained. The mean accurate FRET values for 13 or 23 bp differ between the two FRET-pairs due a difference in Förster radius R_0_. (C) The proposed R_0_-correction allows to convert all accurate FRET values on one common R_0_-axis. (D) ALEX-data for 8, 13, 18, 23, 28 and 33 bp after the 3 correction steps, against background, gamma and R_0_. Error bars were obtained from n = 4 experimental repeats.

To allow a direct comparison of FRET efficiencies with and without PIFE, we suggest the following data analysis procedure (Fig. S2B). **Step 1**: raw data on the level of apparent FRET are corrected for background and spectral crosstalk^49^; this allows to retrieve information on PIFE and possibly on R_1_ if suitable calibration of the PIFE-ruler is available (Fig. S2B). **Step 2**: By subsequent gamma correction, i.e., taking detection and quantum yield differences of donor and acceptor into account^49^ accurate FRET values are obtained. Please note that Cy3 and Cy3-PIFE needs to be treated with a distinct gamma factor. **Step 3**: Finally, a correction for the differing R_0_-values is needed that transforms the relevant FRET populations (Cy3, Cy3-PIFE, Cy3B) on the basis of the same R_0_. For this we use Cy3B as a standard since the quantum yield of the latter is fixed and more or less independent of either the FRET efficiency or of the environment. Comparing two cases with and without PIFE assumes during PIFE only the donor QY, ϕ_D_, is altered (and not dye orientation or other factors) and hence approximate R_0_ in the presence of PIFE as in equation 1:

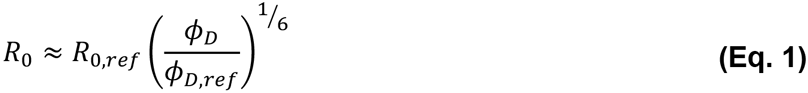

Using the relation between accurate FRET E, interprobe distance *r* and R_0_, we derive the R_0_ corrected FRET efficiency E_R0_ considering the *ℓ*-fold enhancement of the donor-quantum-yield caused by PIFE. This enhancement factor can be obtained directly from the ratio of the two gamma-factors, which are proportional to the quantum yields of Cy3 and Cy3B, assuming constant detection efficiencies and negligible spectral shifts (equation 2):

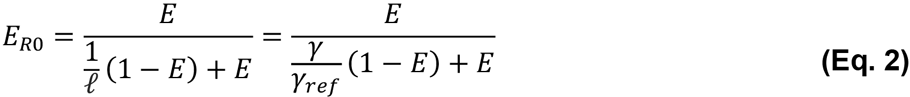

We suggest the reference *γref* to be that of unbound Cy3B-labelled dsDNA. Using the definition of E_R0_ as in Eq. 2, the FRET efficiency is decoupled from *R_0_* changes and related only to distance changes standardized to the *R_0_* value of free Cy3B on dsDNA (See Supplementary Note 1 for a complete derivation of equations 1 and 2 and Fig. S2 for a complete schematic overview of the data analysis).

We tested this procedure by “aligning” the data sets of Cy3-and Cy3B-labelled DNA in combination with FRET acceptor ATTO647N. Using Eq. 2 we normalized the Cy3-data set to that of Cy3B by *R_0_*-correction. As seen in Figure 5C for 13 and 23 bp and all other *R_2_* distances (Fig. 5D, right panel), E_RO_ values of both fluorophore pairs are in excellent agreement validating our data analysis procedure. We note that equation 2 and the information given in Supplementary Note 1 provide a general possibility to account for changes in donor QY even by other photophysical processes such as quenching^95^ (black-hole quencher etc.).

For use of PIFE-FRET in complex biochemical systems, i.e., where correction factors for individual populations may not be accessible, we suggest the use of a simplified data correction procedure. Here, the raw data on the level of apparent FRET are corrected for background and spectral crosstalk^49^. By subsequent gamma correction with the gamma value of Cy3B, we obtain accurate FRET values that are all normalized to the Cy3B-R_0_-axis. This scheme is described in detail in Supplementary Note 2 (eqn. 6-11) and does not require determination of correction factors for individual populations. A comparison of Cy3-and Cy3B-labelled DNA with both analysis methods (see R_0_- correction above) reveals that this method is a valid approximation (Figure S9), but the full correction procedure detailed in Figure 5 remains more accurate.

**PIFE-FRET monitors nucleic acid protein interactions and associated conformational changes**. The data in Figure 5 can only demonstrate the possibility to remove apparent changes in FRET efficiency caused by mere donor QY changes by an “R_0_-correction”. Next, we performed experiments where both PIFE and FRET are altered (and determined) within one experiment due to binding-associated conformational changes of dsDNA. For this we tested the signature of binding of *BamHI* and *EcoRV* in PIFE-FRET at close proximity (R_1_=1 bp) from the donor and at varying R_2_ distances (Fig. 6A). Both restriction enzymes have different binding modes on DNA, i.e., *BamHI* binding should not alter DNA conformation while *EcoRV* induces a 50° kink in the DNA after binding. Hence *BamHI* is expected to show a PIFE effect but preserve FRET after binding, while *EcoRV* should show both PIFE and FRET signal changes. The ruler characteristics of PIFE and FRET, i.e., signal-to base-pair dependence for *BamHI, EcoRV* and Cy3(B)-ATTO647N were described in Figures 4D and 5D.

In these experiments we use the full R_0_-correction and a final normalization to Cy3B-gamma (Figure 6); the provided values of all measurements in Figure 6 can be compared directly. For *BamHI* we expected no changes in the dsDNA conformation but a pronounced PIFE effect after binding (Fig. 6B); the experimentally observed PIFE effect is constant for all observed DNAs (Fig. 6C) and is consistent with the ruler distance for R_1_ of 1 bp. PIFE effect is also only observed when using Cy3 as a donor fluorophore (Fig. 6C). Full data sets including two-dimensional fitting are shown in Figure S10 in the Supplementary Information.

Interestingly, the FRET ruler (distance R_2_) shows a pronounced decrease of *E_R0_* values after binding of *BamHI* (Fig. 6B). Our data allows concluding that this observation is not an artefact of PIFE-FRET nor it is an artefact of our data analysis procedure, but rather a real increase of the donor-acceptor distance, since we observe similar trends when using Cy3B (Figs. 6C/S10), TMR or Atto550 as a donor (data for the latter two not shown).

**Figure 6.**
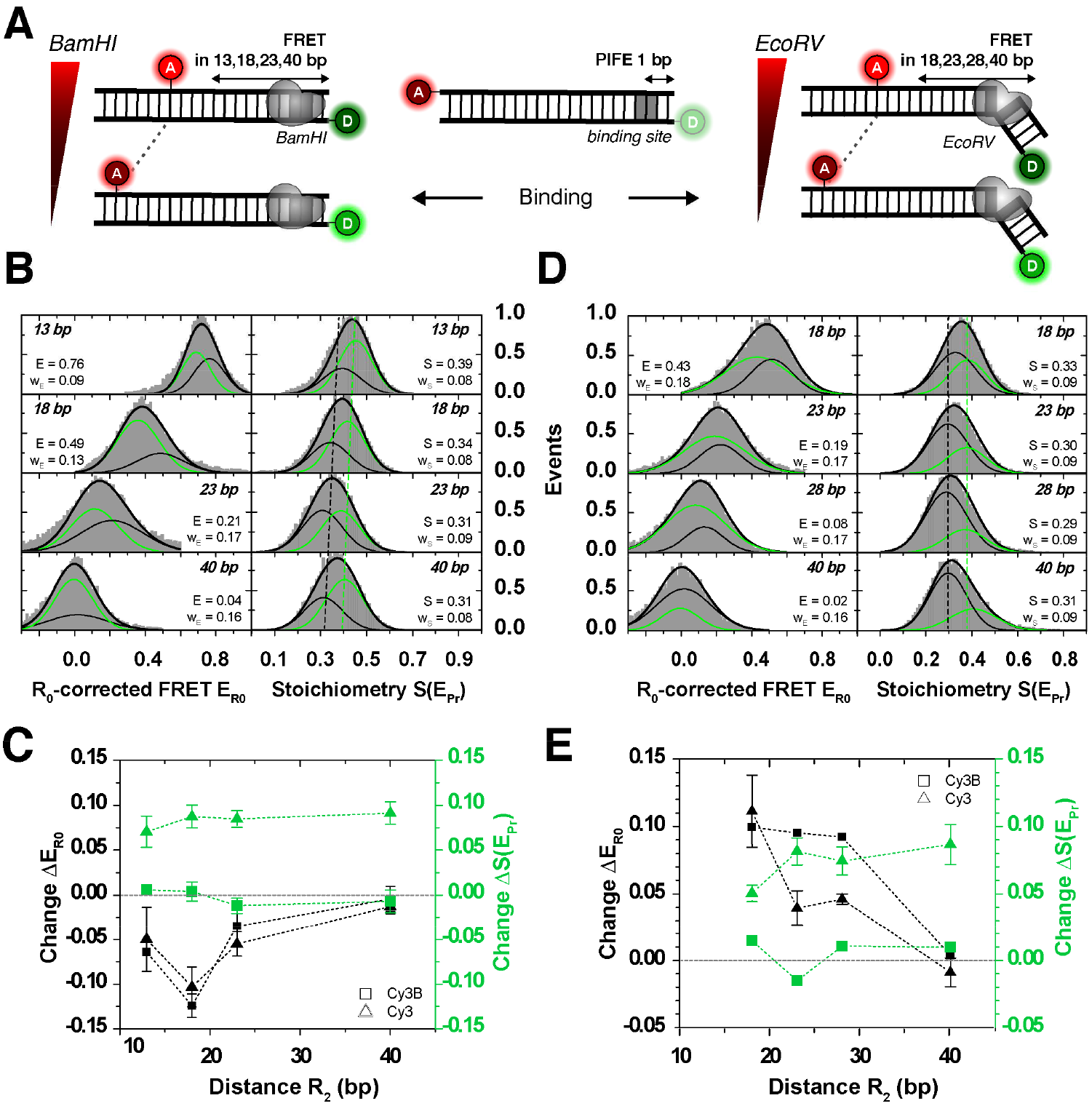
**Distance changes determined via FRET in the presence of PIFE**. FRET between Cy3/ATTO647N attached to a 40 bp-dsDNA is probed simultaneously to PIFE that occurred between Cy3 and different restriction enzymes in R_1_ = 1 bp distance to the donor. (A) Schematic of DNA constructs probing FRET distances in 13, 18, 23 and 40 bp between ATTO647N and Cy3(B) in the presence of *BamHI*, respectively in 18, 23, 28 and 40 bp in the presence of *EcoRV*. While *BamHI* does not interfere with the 3D structure of the DNA, *EcoRV* is reported to introduce a 50° kink. (B) 1D ALEX histograms showing 500 nM *BamHI* bound to 1bp-Cy3-*BamHI*-dsDNA. Binding is detected as constant shift in S between ~ 0.3 (black) and ~ 0.4 (green). *BamHI* introduces a conformational change in the dsDNA seen by smaller FRET values. (C) Differences in Stoichiometry ΔS(E_Pr_) and R_0_-corrected FRET ΔE_R0_ between free and *BamHI*-bound DNA. While binding is observed via PIFE of Cy3 (green, triangle), it is undetected for Cy3B (green, square). The conformational change in FRET is also observed for Cy3B (black, square). (D) 1D ALEX histograms showing 50 nM *EcoRV* bound to 1bp-Cy3-*EcoRV*-dsDNA. Binding is observed as a constant shift in S from 0.29 to 0.38. EcoRV is reported to kink dsDNA, which is observed by increased FRET values. (E) Differences in Stoichiometry ΔS(E_Pr_) and R_0_-corrected FRET ΔE_R0_ between free and *EcoRV*-bound DNA labelled with Cy3 (triangle) or Cy3B (square). While distance changes in R_2_ for different FRET distances are observed for both donor fluorophores (black panel), binding is only observed via PIFE to Cy3 (green, triangle) and undetected for Cy3B (green, square). Error bars were obtained from at least n = 2 experimental repeats. Full data sets and controls provided in Figure S10.

The current understanding of *BamHI*-interactions with dsDNA suggests that there should be only minor structural changes in dsDNA after binding^91^. We hence hypothesize that the FRET decrease (Figs. 6B/C) could correspond to a reduction of the accessible volume of Cy3(B) due to steric restriction of the dyes or to changes of orientation factor *κ*^2^ due to changes of fluorophore interaction with dsDNA due to the BamHI protein Steady-state anisotropy values were determined for external Cy3 (r_D_ = 0.231 ± 0.026), internal Cy3 (r_D_ = 0.175 ± 0.036), external Cy3B (r_D_ = 0.191 ± 0.007) and internal ATTO647 (r_A_ = 0.110 ± 0.014). In presence of BamHI the anisotropy values showed a slight increase for all samples: external Cy3 (r_D_ = 0.259 ± 0.033) and internal Cy3 (r_D_ = 0.205 ± 0.044), Cy3B (r_D_ = 0.235 ± 0.032), and ATTO647N (r_A_ = 0.149 ± 0.019). While both Cy3 and Cy3B show high values that indicate stacking or groove binding^97^, ATTO647N has significantly lower anisotropy values. A published theory^98^ can be used in combination with the determined anisotropy to yield the associated distance uncertainty, which we provide Supplementary Table 4. It amounts maximally to 25% uncertainty of the final distance determined.

Additional control experiments with the DNA-sliding T7 polymerase gpt/trx (gp5/trx^88–90^, Figs. S3/11) that binds non-specifically to dsDNA confirms our data analysis procedure (R_0_-correction, eqn. 1+2): T7 polymerase binding on dsDNA results in pronounced PIFE effects but constant ER_0_ values (Fig. S11) verifying that the observed FRET changes for BamHI binding are real and not caused by convolution of PIFE and FRET effects.

Performing similar experiments with *EcoRV* shows results that agree with structural predictions and reveal the full power of PIFE-FRET. A PIFE-effect consistent with 1 bp separation is found for all DNAs (Figs. 6D/E), which can only be observed for use of Cy3 as a donor fluorophore (Fig. 6E). As expected from the well-established kinking of *EcoRV* of dsDNA upon binding, the FRET ruler suggests a decrease in the donor-acceptor distance consistent upon binding and related DNA-kinking (Figs. 6D/E).

**PIFE-FRET maps initial transcription of E. coli RNA polymerase**. Finally, we used PIFE-FRET to study the *E. coli* RNA polymerase (RNAP) promoter DNA complex that still lacks full structural characterization due to its dynamic nature especially during transcription initiation. In an *in vitro* transcription assay of RNAP holoenzyme (core and σ^70^ transcription factor) and lacCONS+20A model promoter, the enzyme melts and forms a ~13 bp DNA transcription bubble (registers −11 to +2) around the transcription start site at register +1 to form a transcription-competent RNAP-promoter open complex (*RP_O_*). Subsequently, different subsets of nucleotide triphosphates (NTPs) were added in order to induce permanent abortive initiation pre-defined by the promoter sequence.^99^ Through this procedure, different RNAP-promoter initially transcribed complex states (RP_ITC≤i,_ i — the maximal length of transcribed RNA, Figure S4E)^100,101^ are visited at which the DNA bubble is scrunched by RNAP^102–104^ allowing downstream DNA bases to melt and serve as reading template for polymerization of RNA.

To monitor the conformation of the DNA bubble in conjunction with its interaction with RNAP, we designed two PIFE-FRET assays with labelling of acceptor ATTO647N at −15 in the template (t) strand and the donor Cy3(B) at position +1/+3 at the nontemplate (nt) strand (Figure 7A). The ALEX data was analysed according to the simplified correction model outlined in Supplementary Note 2.

**Figure 7.**
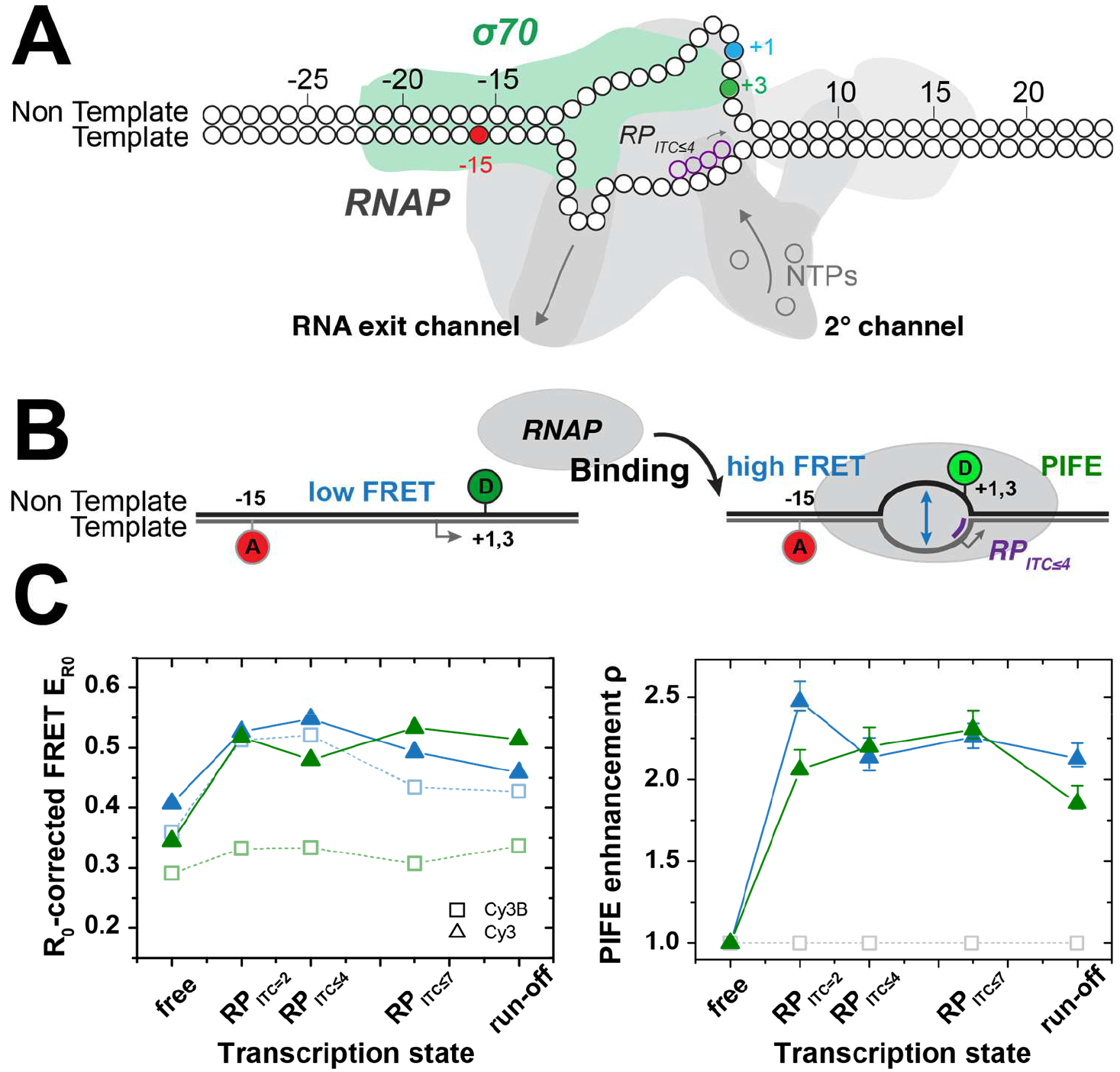
**Mapping the proximity of the scrunched transcription bubble with RNAP**. (A) Labelling scheme. The acceptor ATTO647N is positioned on the template strand at register (-15). The donor Cy3(B) is placed on the non-template strand at register (+1) (cyan) or (+3) (green). The dsDNA bound to the RNAP is depicted in an illustration of the transcription initiation state RP_ITC≤4_ (B) Probing the transcription bubble formation during RNA transcription initiation using PIFE-FRET. FRET reports on the DNA opening. The increase in PIFE reports on binding of RNAP and conformational changes of the DNA bubble. (C) Probing the scrunched transcription bubble for RNA transcription initiation in *E. coli* via PIFE-FRET formed at a lacCONS+20A dsDNA promoter^105,106^. The R_0_-corrected FRET values (left panel) and PIFE enhancement *ρ* (right panel) as provided for the registers −15/+1 (blue) and −15/+3(green). These results were achieved by a global fit of the data to the model provided by ref.^52^ (see Fig. S12).

For RNAP binding to promoter DNA, both +1/+3 donor *nt* registers report on bubble opening via a FRET increase from free DNA to RP_ITC=2_. Scrunching is most pronounced in NTP-starved initiation states, RP_ITC≤4_ and RP_ITC≤7_ and having no full return of the bubble region to dsDNA in the run-off product shows RNAP is unable to run-off the labelled promoter sequence. These observations are in line with previous studies of initial transcription of E. coli RNAP. However, our data reveal the fine details of DNA scrunching when comparing how the bubble conformation changes when probes in reference to different *nt* registers. More specifically, register +1, which precedes register +3 in being part of the transcriptional bubble at an earlier initiation stage, shows an increase in FRET until reaching RP_ITC≤4_ then a decrease in FRET moving from RP_ITC≤4_ to RP_ITC≤7_, while register +3 seems to show this trend only starting at RP_ITC≤7_.

The PIFE readout shows S(E) values of free promoter DNA that change upon transition to initial transcribing complexes. In order to decouple PIFE increases in S from changes in S caused by an increase in E with no PIFE, the S(E) values were globally fitted to the model suggested in^52^ (best fit results are shown as isolines in left panels). The absolute PIFE parameter values, which report on the fold-decrease in Cy3 cis/trans isomerization mobility, report on the complex changes in the proximity of the transcriptional bubble to RNAP most probably caused by scrunching. Interestingly Cy3 on *nt* register +1 shows a maximal PIFE effect in RP_ITC=2_, then reduces upon scrunching in states RP_ITC≤4_ and RP_ITC≤7_. This reduction in the PIFE effect is accompanied by an increase in the PIFE effect for *nt* register +3 that is introduced to the bubble only after scrunching. (Figs. 7B/C). Regarding PIFE, the increase in peak stoichiometry for Cy3 is a first direct observation of RNAP binding in the vicinity of Cy3 close to position (+1/+3) of the *nt* strand.

These combined trends can possibly be interpreted as follows: during abortive initiation of transcription, the position of Cy3 in the *nt* strand as well as the structure of the DNA bubble vary relative to RNAP due to scrunching. Lately Winkelman et al.^68^ have shown how, in RP_ITC_ states, *nt* registers (-2) - (+1) are extruded out of the surface of RNAP, by using cross-linking methods. The reduction in the PIFE with Cy3 labelling this specific position in the *nt* strand may indicate that this NTP-starved RP_ITC_ state characterized by extrusion of the *nt* strand out of the RNAP surface exists also free in solution and is not only an outcome of possible cross-linking artefacts. In addition, these finding are strongly supported by the similar trends in the changes the conformation of the transcription bubble undergoes, as judged by FRET in reference to registers +1 and +3 in the *nt* strand.

The effect observed through the FRET ruler can be interpreted as directly responding to the DNA scrunching, which shortens the distance between the downstream edge of the transcription bubble and bases upstream to it. However, how can the decrease in FRET efficiency in RP_ITC≤7_ be explained? Lerner and Chung have lately revealed the existence of a state in which transcription pauses for long periods of time and the nascent RNA backtracks by 1-2 bases into the secondary channel of RNAP in RP_ITC_≤7.^99^ One outcome is the decrease in bubble size and scrunching as confirmed by magnetic tweezer experiments.^99^ Following the FRET results, a reduction in the bubble size can be inferred from the results using the *nt* register +1.^99^ Although a reduction in the transcriptional bubble size and in scrunching is expected, how may it be different in the paused-backtracked state from the perspective of the bubble conformation? The simultaneous measurement of this effect through both FRET and PIFE offers an intriguing potential alternative route for the release of the strain built up by scrunching through the extrusion of the nontemplate strand out of the surface of RNAP.

Overall, these findings illustrate that PIFE-FRET allows probing of global conformational changes using FRET as well as changes in positioning of proteins along DNA, also in states where FRET changes are only subtle. The results shown here (Fig. 7) are the first report of a” “reagent-free optical footprint” of RNAP performing DNA scrunching in initiation. Scanning Cy3 attachment to all relevant template and nontemplate registers will yield a full picture of even the dynamic aspects of the RNAP footprint bound to DNA in transcription initiation.

## DISCUSSION

To date there are only few (single-molecule) assays allowing the simultaneous observation of both protein binding and the conformational changes associated with the binding event. One way is multi-colour ALEX which utilizes a FRET cascade of more than two fluorophore probes^50, 107,108^. Although a powerful technique, it requires dye labelling of both nucleic acid and protein and sophisticated analysis that go even beyond the procedures introduced here.

In this work, we realized a novel combination of protein-induced fluorescence enhancement (PIFE) and Förster resonance energy transfer (FRET). Our proposed technique allows for mechanistic investigations of protein-nucleic acid interactions with diffusion-based confocal microscopy (demonstrate in this work) or surface-immobilized molecules^60–62,109^ without labelling of the protein of interest. This approach is compatible for single-molecule studies of weak (μM) interactions. After accounting for PIFE effects coupled to FRET, one can utilize PIFE-FRET to probe binding, global conformational changes in the Förster distance scales (3-10 nm) and local conformational changes on shorter distance scale (< 3 nm) and hence use the assay as a multi-scale quantitative ruler. Nevertheless, whereas the FRET dependence on distance between probes is well-established and general, the distance dependence of the PIFE effect requires careful characterization for each system under investigation, as it highly depends on the spatial topology of the binding surface, which brings about different steric hindrance effects for different binding modes of DNA with different proteins. In case of RNAP, we envision systematic experiments of RNAP e.g., free promoter DNA and *RP_O_* with the Cy3 dye labeling different registers on both the nontemplate and template strand, i.e., in the transcription bubble as well as at downstream positions. The results shown in Figure 7 re-affirm the existence of a scrunched DNA state^68^, in which register +1 of the nontemplate strand is extruded out of the surface of RNAP in initially transcribing complex, while the size of the scrunching-enlarged transcription bubble decreases. This was found by use of PIFE-FRET without the use of chemical cross-linking agents. We therefore name these types of experiments by the name ‘reagent-free optical footprinting’ and envision the technique will allow the characterization of (dynamic) conformational states, such as RP_ITC≤i_ states, that may be biased by the cross-linking procedure and are hard to capture using X-Ray crystallography.

As shown in Figure S3, the PIFE technique and also PIFE-FRET is not exclusive for cyanine fluorophores and Cy3 but other fluorescence quenching^95^ (or enhancing) mechanisms can be used. An alternative mechanism is photo-induced electron transfer (PET) whereby tryptophan or guanosine may be used as specific fluorescence quenchers^110–112^. A FRET-PET hybrid technique has been recently introduced^113^ and might, in principle, be coupled with PIFE. Nevertheless, this technique requires usage of specific fluorophores susceptible for PET by Trp, and also a Trp carrying protein with a well-characterized positioning for the PET to occur. With PIFE-sensitive fluorophores, the restriction of its steric freedom by a nearby bound protein alone is enough to induce the PIFE-effect. In addition, the dependence of a PET effect with respect to the dye-quencher distance is at very short molecular separations and often even binary (ON/OFF). In this extreme case PET would only report on molecular contact but not act as a ruler. PIFE, however, can report (under some conditions) continuously on distances up to 3 bp separation between dye and the surface of a bound protein, hence of more general applicability for probing complex biochemical processes.

In general all possible specific interactions of both dyes with the protein (hydrophobic effects, PET via Trp/Tyr-residues, singlet quenching etc.) have to be considered since these alter the observed “apparent” PIFE effect. The effect of steric hindrance serves as a lower boundary to the PIFE effect^52^ and Cy3 at some bases (3 and 5 bp from interaction with BamHI) can have a larger PIFE effects than expected just from steric hindrance, which can only be explained by additional specific interactions. Thus, for a completely unknown biochemical system or conformation, it would be hard to fully deconvolve the PIFE-distance from FRET by only relying on ratiometric measure of donor and acceptor intensities using μsALEX. Ultimately, we think that an optimized PIFE-FRET assay would include donor-and acceptor state-selective lifetimes to disentangle all possible contributions from the experimental signature. Since an acceptor-based PIFE-FRET assay also obliviates the need for an R_0_-correction, lifetime-and acceptor-based experiments will likely be the future of PIFE-FRET. This requires, however, acceptor optimal PIFE-fluorophores in the red spectral region that serve as FRET acceptor. Since PIFE depends on the 1D polymethine linkage, fluorophores such as Cy5 have many more possibilities for bond rotations compared to Cy3, complicating the competition between fluorescence and cis-trans isomerization, thus reducing possible PIFE-effects^52^ (see Suppl. Figure S13 for a comparison of Cy3 and Cy5 PIFE effects with BamHI). An ideal strategy would be to increase the conjugated chromophore system not via extension of the polymethine chain as e.g., from Cy3 to Cy3.5. Another possible solution would be the use of a blue/green-based PIFE-FRET assay, e.g., with Alexa488/ATTO488 (donor) and Cy3 (acceptor), which likely suffers from increased spectral crosstalk and limitations in the photophysical properties of blue fluorophores.

To finally relate the distance dependencies of the PIFE-ruler present here to published smPIFE work, the fold increase in Cy3 fluorescence QY due to the PIFE effect is needed, which is only indirectly available in ALEX. In a forthcoming paper^52^, we provide the needed photophysical framework for ALEX-based PIFE-FRET to demonstrate its ability to obtain fully quantitate experimental results from ALEX that are directly comparable with published smPIFE studies. The nature of the model described in ref.^52^ might also allow to include other contributions that change either donor or acceptor properties by (static or dynamic) singlet quenching.

## MATERIAL AND METHODS

**DNA, Proteins and Reagents**. Unless otherwise stated, reagents of luminescent grade were used as received. Amino-modified and fluorophore-labelled oligonucleotides were used as received (IBA, Germany). DNA single strands were annealed using the following protocol: A 5-50 μL of a 1 μM solution of two complementary single-stranded DNAs (ssDNA) was heated to 98 °C for 4 minutes and cooled down to 4 °C with a rate of 1 °C/min in annealing buffer (500 mM sodium chloride, 20 mM TRIS-HCl, and 1 mM EDTA at pH = 8).

Different sets of complementary DNA-oligonucleotides were used (Fig. S4). <u>Set 1</u>: The first scaffold uses two complementary 45-mers carrying the donors (Cy3, Cy3B, TMR or Alexa555) at the 5’-end of the top-strand (Fig. S4A). The acceptor (ATTO647N or Cy5) was attached in 8, 13, 18, 23, 28 or 33 bp separations to the donor fluorophore. The DNAs are referred to as e.g., 33bp-Cy3/ATTO647N for a sample with 33 bp separation between Cy3 on the top-strand and ATTO647N on the bottom-strand (Fig. S4A). Non-specific binding of T7 DNA polymerase gp5/thioredoxin via PIFE was investigated using 18/23/28/33bp-Cy3/ATTO647N as well as 33bp-(Cy3/Cy3b/AF555/TMR)/ATTO647N and 33bp-TMR/Cy5. T7 DNA polymerase gp5 was expressed and purified in a 1:1 complex with thioredoxin in E. coli^114^. These samples were provided by the labs of van Oijen and Richardson.^114^. <u>Set 2</u>: To study the distance dependence of PIFE in absence of FRET we used DNAs comprising of 40-mers carrying Cy3(B) and ATTO647N (Fig. S4B). These DNAs separate both dyes by 40 bp which prohibits FRET-interactions due to large separation >10 nm. The DNAs carry palindromic sequences for two different restriction enzymes *BamHI* and *EcoRV* at 1,2,3,5 and 7 bp distance with respect to Cy3(B) and are termed 1bp-Cy3(B)-(#bp)- *BamHI*-40-ATTO647N for a dsDNA with 40 bp separation in FRET and #bp separation between the donor and *BamHI* inducing PIFE. DNA sequences and positioning of *BamHI* binding sites were adapted from ref.^16^; those for EcoRV were derived from 1 bp-PIFE-*BamHI*-DNA^16^. <u>Set 3</u>: To study the distance dependence of PIFE in presence of FRET, complementary 40-mer oligonucleotides carrying the donors (Cy3 and Cy3B) at the 5’-end of the top-strand and palindromic binding sequence for *BamHI* and *EcoRV* in 1 bp distance from the donor were employed (Fig. S4C). The acceptor (ATTO647N) was attached in 13,18,23 and 40 bp (*BamHI*) respectively in 18,23,28 and 40 bp (*EcoRV*) distance to the donor fluorophore. The DNAs are termed (analogue to Set3) 1 bp-Cy3(B)-1bp-*BamHI*-(#bp)-ATTO647N for a dsDNA with 1 bp separation in PIFE and #bp separation between the donor and the acceptor. DNA sequences and positioning for *BamHI* and *EcoRV* were derived from 1bp-PIFE-*BamHI*-DNA^16^. <u>Set 4</u>: To check the influence of internal and external labelling (Fig. S7) we attached Cy3 to the 3^rd^ base pair in the top strand of 5bp-PIFE-*BamHI*-DNA and ATTO647N at the 5’end of the bottom strand (Fig. S4D). We termed it 3bp-Cy3(B)-1bp-*BamHI*-40-ATTO647N. *BamHI* and *EcoRV* were used as received (NEB/Bioké, The Netherlands). <u>Set 5</u>: PIFE-FRET was employed to study the interaction between *E.Coli* RNA polymerase and promoter dsDNA during transcription initiation. In these experiments Oligos (Fig. S4E-F) having the lacCONS+20A sequence^106^. The template and nontemplate strands, were labeled with ATTO647N and Cy3(B) at different promoter registers as indicated in Figure S4 (IBA oligos, GmbH). RNAP holoenzyme was supplied by NEB, Ipswich, MA, USA, M0551S. High-purity ribonucleotide triphosphates (NTPs) (GE Healthcare, Little Chalfont, Buckinghamshire, UK) as well as Adenylyl(3′-5′) adenosine (ApA; Ribomed, Carlsbad, CA, USA) were used in all transcription reactions at 100 μM each.

ALEX-experiments were carried out at 25-50 pM of dsDNA at room temperature (22°C). For experiments on dsDNA only (Fig. S4A) or in combination with gp5/trx, an imaging buffer based on 50 mM TRIS-HCl, 200 mM potassium chloride at pH 7.4 was applied. 1 mM Trolox^115,116^ and 10mM MEA were added to the buffer for photostabilization as reported in ref.^117^. Experiments with *BamHI* were carried out in 50 mM TRIS-HCl, 100 mM sodium chloride, 10 mM CaCl2 and 0.1 mM EDTA at pH 7.4 in the presence of 143 mM bME. Experiments with *EcoRV* were carried out in 50 mM TRIS-HCl, 100 mM sodium chloride, 10mM CaCl2 and 0.1 mM EDTA at pH 7.4. All binding experiments with *BamHI* and *EcoRV* were performed in the presence of calcium chloride, to prevent enzymatic activity^91–93^ and the formation of aggregates^94^.

**RNA polymerase transcription assays**. 10 μl of pre-solutions were prepared by 180 nM RNA Polymerase (RNAP) holoenzyme in TB buffer (50 mM Tris-HCl, pH 8, 100 mM KCl, 10 mM MgCl2, 1 mM DTT, 100 μg/ml BSA, and 5% glycerol). Solutions were incubated 20 min at 30°C, then 0.6μl of 1μM promoter DNA was added, and samples were further incubated 30 min at 37°C. 1 μl of 100 mg/ml Heparin-Sepharose suspension (GE Healthcare, Inc.) was added with 20 μl of KG7 buffer (40 mM HEPES-NaOH, pH 7, 10 mM MgCl_2_, 1 mM DTT, 1 mM MEA, and 100 μg/ml BSA), then incubate 1 min at 37°C, to eliminate free RNAPs as well as RNAP binding promoter non-specifically. After 1 min incubation, samples were centrifuged using a table top centrifuge, and 15 μl of supernatants were transferred to tubes containing 15 μl of KG7 buffer incubate 30 min at 37°C to make RP_O_ solutions. In order to make each transcription initial complexes, 4 μl of RP_O_ solutions are transferred into 16 μl of solutions containing 0.625 mM A_p_A (for RP_ITC_=2), 0.625 mM A_p_A + 0.625 mM UTP (for RP_ITC_≤4), 0.625 mM A_p_A + 0.625mM UTP + 0.625 mM GTP (for RP_ITC_≤7), 0.625 mM ATP+0.625 mM UTP+0.625 mM GTP (for RD_E_=11)^106^, or 0.625 mM of all NTPs (for Runoff)^106^, in KG7 buffer, then incubate 30 min at 37°C.

**Fluorescence and anisotropy measurements**. Fluorescence spectra and anisotropy ^118^ values *R* were derived on a standard scanning spectrofluorometer (Jasco FP-8300; 20nm exc. and em. Bandwidth; 8 sec integration time) and calculated at the emission maxima of the fluorophores (for Cy3B, λ_ex_ = 532 nm and λ_em_ = 570 nm; for ATTO647N, λ_ex_ = 640 nm and λ_em_ = 660 nm), according to the relationship *R* = (I_VV_ - GI_VH_)/(I_VV_+2GI_VH_). I_VV_ and I_VH_ describe the emission components relative to the vertical (V) or horizontal (H) orientation of the excitation and emission polarizer. The sensitivity of the spectrometer for different polarizations was corrected using horizontal excitation to obtain G = I_HV_ / I_HH_.

**Fluorescence lifetime measurements**. Fluorescence lifetimes were determined using time-correlated single-photon counting with a home-built confocal microscope described in ref.^119^. Fitting of the decay functions was done with a mono-(Cy3B, ATTO647N) or double-exponential function (Cy3) taking the instrumental response into account. Values reported in this section and Table 1 are given with an error of 5%. The data was processed via a custom data evaluation program^120^ written in MATLAB (2013b, MathWorks Inc., Natick, MA). The procedure yielded a bi-exponential decay of 1.6 and 0.4 ns for Cy3, and mono-exponential decays of 2.29 ns for Cy3B and 4.24 ns for ATTO647N on a 40-mer dsDNA. The presence of *BamHI* alters the lifetimes to 1.75 ns and 0.4 ns on average (Cy3), 2.22 ns (Cy3B) and 4.29 ns (ATTO647N). On a 45-mer DNA lifetimes of Cy3 (1.18 ns, bi-exponential average), Cy3B (2.30 ns), TMR (3.23 ns), Alexa555 (1.64 ns), ATTO647N (4.17 ns) and Cy5 (1.38 ns) were determined. Addition and non-specific binding of gp5/trx alters their lifetimes as follows: Cy3 (1.69 ns, biexponential average), Cy3B (2.5 ns), TMR (2.98 ns), ATTO647N (4.23 ns) and Cy5 (1.70 ns).

*ALEX-Spectroscopy and data analysis*. For single-molecule experiments custom-built confocal microscopes for μs-ALEX described in^121,122^ were used as schematically shown in Figure S2. Shortly, the alternation period was set to 50 μs, and the excitation intensity to 60 μW at 532 nm and 25 μW at 640 nm. A 60x objective with NA=1.35 (Olympus, UPLSAPO 60XO) was used. Laser excitation was focused to a diffraction limited spot 20 μm into the solution. Fluorescence emission was collected, filtered against background (using a 50-μm pinhole and bandpass filters) and detected with two avalanche photodiode detectors (τ-spad, Picoquant, Germany). After data acquisition, fluorescence photons arriving at the two detection channels (donor detection channel: D_em_; acceptor detection channel: A_em_) were assigned to either donor-or acceptor-based excitation on their photon arrival time as described previously.^67,85^ From this, three photon streams were extracted from the data corresponding to donor-based donor emission F(DD), donor-based acceptor emission F(DA) and acceptor-based acceptor emission F(AA; Fig. S2A).

During diffusion (Fig. S2B), fluorophore stoichiometries S and apparent FRET efficiencies E^*^ were calculated for each fluorescent burst above a certain threshold yielding a two-dimensional histogram.^67,85^ Uncorrected FRET efficiency E^*^ monitors the proximity between the two fluorophores and is calculated according to:

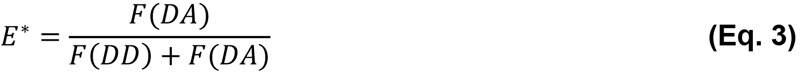

S is defined as the ratio between the overall green fluorescence intensity over the stotal green and red fluorescence intensity and describes the ratio of donor-to-acceptor fluorophores in the sample S:

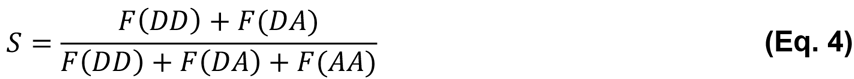

Using published procedures to identify bursts corresponding to single molecules^123^, we obtained bursts characterized by three parameters (M, T, and L). A fluorescent signal is considered a burst provided it meets the following criteria: a total of L photons, having M neighbouring photons within a time interval of T microseconds. For all data presented in this study, a dual colour burst search^123,124^ using parameters M = 15, T = 500 μs and L = 25 was applied; additional thresholding removed spurious changes in fluorescence intensity and selected for intense single-molecule bursts (all photons > 100 photons unless otherwise mentioned). Binning the detected bursts into a 2D E^*^/S histogram where sub-populations are separated according to their S-values. E^*^- and S-distributions were fitted using a Gaussian function, yielding the mean values *μ_i_* of the distribution and an associated standard deviations *w_i_*. Experimental values for E^*^ and S were corrected for background, spectral crosstalk (proximity ratio E_PR_) and gamma factor resulting in histograms of accurate FRET E and corrected S according to published procedures^49^.

**Data analysis to retrieve distance Ri (PIFE-ruler)**. All data were corrected against background and spectral crosstalk to yield E_PR_ and S(E_PR_). To determine the induced enhancement introduced by the change in Cy3 cis/trans isomerization monility either by viscosity (glycerol; Fig. 3) or by steric hindrance (caused by a binding protein close-by), the mean value of stoichiometry S(E_Pr_) of the free DNA was determined via 2D-Gaussian fitting of the 2D E_Pr_-S(E_Pr_)-histogram. DNA in the presence of a DNA-binding protein was fitted with two independent 2D Gaussian population, where one population - the unbound species - was approximated with constant values obtained for the free DNA species before. The observed PIFE enhancement was represented as difference in Stoichiometry S(E_Pr_). The PIFE enhancement factor ρ that reports on the PIFE effect decoupled from S changes caused by pure E changes (without PIFE) was retrieved from an advanced model as described in ref.^52^. In brief, this model allows for the retrieval of the amount by which the excited-state *trans/cis* isomerization of Cy3 is slowed down (the fold decrease in *cis/trans* isomerization mobility).

**Data analysis to retrieve distance R_2_ (FRET-ruler) in the presence of PIFE**. All data was corrected against background and spectral crosstalk. At <u>first</u> the γ_Cy3(B)_ for all free DNAs was determined and free DNA’s data was corrected until accurate FRET E. For this, all data needs to be corrected against background and spectral crosstalk. For both FRET pairs the individual gamma factors, γ_Cy3(B)_ were determined, and each population was corrected with it obtaining accurate FRET E. In a <u>second</u> step, the gamma factor for the protein bound species γ_Cy3(B)_/protein is determined, and each population within the data set is corrected with its own individual γ_Cy3(B)_/protein and γ_Cy3(B)_/free. This is achieved by assigning each burst of the uncorrected data at the beginning to either the free or bound DNA subpopulation (see next paragraph). This is followed by a selective accurate FRET correction for each subpopulation. After this correction step all determined R_0_-corrected FRET values for the free and bound Cy3-dsDNA are converted onto the R_0_-axis of the environmentally insensitive Cy3B by applying Eq. 2 burst-wise. The mean R_0_-corrected FRET value E_R0_ is determined by 2D-Gaussian fitting of the E_R0_-S-histogram. FRET values of converted-Cy3 and Cy3B should be identical within errors at this correction stage. As convention, we transformed all presented R_0_-corrected FRET values onto the unaltered R_0_-axis of the free Cy3B-labeled DNA in this manuscript.

**Population assignment**. In order to correct individual populations with different correction factors, as gamma factors, within one 2D ALEX histogram, every burst needs to be assigned to a particular population. This can be achieved via cluster analysis methods or probability distribution analysis^125^. In our implementation, every population in the uncorrected 2D histogram is first fitted with a covariant bivariate Gaussian function

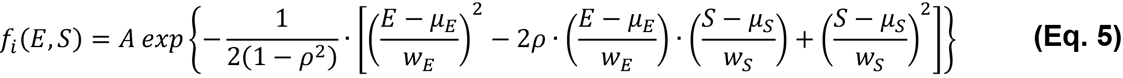
where the population is described by an amplitude *A*, its mean values *μ_i_* and standard deviations *w_i_* in FRET E^*^ and Stoichiometry S. *ρ* denotes the correlation matrix between E^*^ and S. We express the probability *p* that a given burst in the 2D histogram belongs to population *i* by

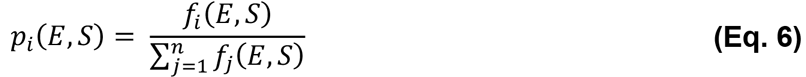

For every bin in the 2D histogram, the algorithm calculates the number of bursts *n_i_* belonging to population *i* by *n_i_ = n.p_i_(E,S)*, where *n* is the number of burst in one bin. E and S are taken to be the bin centre. The corresponding bursts are assigned to a particular population 2 and kept through out the data analysis process.

## AUTHOR CONTRIBUTIONS

T.C. conceived the study. E.P., E.L., S.W., J.H. and T.C designed research. E.P., E.L., F.H., and S.C. performed research. E.L., J.H. and M.R. provided new analytical tools. E.P., E.L., M.R. and T.C. analysed data. The manuscript was written through contributions of all authors. All authors have given approval to the final version of the manuscript.

## NOTES

The authors declare no competing financial interests.

## ACKNOWLEDGMENT

The authors are grateful to A.M. van Oijen for generous and enduring support of this study. We thank A.M. van Oijen and C. Richardson for the kind gift of T7 DNA polymerase 5 (gp5/trx), A. Aminian Jazi for support, G. Gkouridis and M.J. de Boer for fruitful discussions and advise.

